# Obesity-associated MRAP2 variants impair multiple MC4R-mediated signaling pathways

**DOI:** 10.1101/2024.11.19.622403

**Authors:** Rachael A. Wyatt, Aqfan Jamaluddin, Vinesh Mistry, Caitlin Quinn, Caroline M. Gorvin

## Abstract

The melanocortin-4 receptor (MC4R) is a G protein-coupled receptor expressed at hypothalamic neurons that has an important role in appetite suppression and food intake. Mutations in MC4R are the most common cause of monogenic obesity and can affect multiple signaling pathways including Gs-cAMP, Gq, ERK1/2, β-arrestin recruitment, internalization and cell surface expression. The melanocortin-2 receptor accessory protein 2 (MRAP2), is a single-pass transmembrane protein that interacts with and regulates signaling by MC4R. Variants in MRAP2 have also been identified in overweight and obese individuals. However, functional studies that have only measured the effect of MRAP2 variants on MC4R-mediated cAMP signaling have produced inconsistent findings and most do not reduce MC4R function. Here we investigated the effect of twelve of these previously reported MRAP2 variants and showed that all variants that have been identified in overweight or obese individuals impair MC4R function. When expressed at equal concentrations, seven MRAP2 variants impaired MC4R-mediated cAMP signaling, while nine variants impaired IP3 signaling. Four mutations in the MRAP2 C-terminus affected internalization. MRAP2 variants had no effect on total or cell surface expression of either the MRAP2 or MC4R proteins. Structural models predicted that MRAP2 interacts with MC4R transmembrane helices 5 and 6, and mutations in two MRAP2 residues in putative contact sites impaired the ability of MRAP2 to facilitate MC4R signaling. In summary, our studies demonstrate that human MRAP2 variants associated with obesity impair multiple MC4R signaling pathways and that both Gs-cAMP and Gq-IP3 pathways should be assessed to determine variant pathogenicity.

## Introduction

Food intake and body weight are centrally regulated by the leptin-melanocortin system. In the fed state, leptin is secreted from adipocytes and acts upon the leptin receptor expressed on neurons within the arcuate nucleus of the hypothalamus that secrete proopiomelanocortin (POMC). POMC is processed post-translationally by enzymes including the prohormone convertase 1 (PCSK1) to the melanocortin peptides (α-β-γ-melanocyte stimulating hormone (MSH)). These peptides are agonists for the G protein-coupled receptor melanocortin-4 receptor (MC4R) expressed at neurons of the paraventricular nucleus (PVN), which suppresses appetite and food intake. Mutations within several genes of the leptin-melanocortin pathway cause monogenic early-onset severe obesity, often associated with hyperphagia, hyperinsulinaemia and in some cases developmental delay. These include genes that regulate neuronal differentiation and development such as the transcription factor single-minded 1 (SIM1), the neurotrophin BDNF and the tropomyosin receptor kinase B (TrkB) by which BDNF signals (1, 2). Genes are also mutated within the leptin pathway including: inactivating homozygous or compound heterozygous variants in leptin or its receptor (LEPR) (3, 4), large deletions in Src-homology-2 B-adaptor protein-1 (SH2B1), by which LEPR signals (5), heterozygous mutations in the steroid receptor coactivator-1 (SRC-1) (6), and deleterious mutations in the nuclear plekstrin homology domain interacting protein (PHIP) which both enhance POMC transcription (7). Within the melanocortin pathway homozygous or compound heterozygous mutations have been identified in PCSK1(8), heterozygous or homozygous loss-of-function mutations in MC4R are the most common cause of severe monogenic obesity (9, 10), while heterozygous inactivating mutations in the GNAS gene encoding the stimulatory G alpha protein (Gαs) by which MC4R signals also cause obesity (11). Heterozygous mutations have been identified by several groups in the melanocortin 2 receptor accessory protein 2 (MRAP2), a single-pass transmembrane protein that has been shown to regulate signaling by MC4R.

MRAP2 was originally identified as a homolog to MRAP1, a transmembrane protein that is essential for the cell surface expression and signaling of melanocortin-2 receptor (MC2R), which regulates adrenal development and steroidogenesis (12). As MRAP2 is highly expressed in the brain it was hypothesised that it may regulate MC4R function, and mouse models depleted of MRAP2 have extreme obesity, increased fat mass and visceral adiposity, analogous to MC4R knockout mice (13, 14). Double knockouts of MRAP2 and MC4R further demonstrated that MRAP2 facilitates the action of MC4R, but that there are also MC4R-independent mechanisms (15). Unlike MRAP1, MRAP2 is not essential for MC4R cell surface expression, although some studies have suggested it may modify receptor expression (12, 16–19). Most studies agree that MRAP2 directly interacts with MC4R (12, 16) and enhances cAMP signaling (15, 20), while a recent preprint has indicated that MRAP2 may impair β-arrestin recruitment to MC4R and facilitate coupling to Gs and Gq proteins (20).

Several studies have described heterozygous MRAP2 variants in association with human obesity. In the first study, one nonsense and three missense variants were identified in individuals from well characterised cohorts of patients with obesity (15) (Table 1). Subsequently, additional missense variants were identified in individuals with Prader-Willi or Prader-Willi-like Syndrome (21). The first study to functionally assess three MRAP2 variants identified in children with extreme obesity, showed only one variant affected MC4R-generated cAMP responses (22), although only a single concentration (1μM) of αMSH was used in an endpoint assay (AlphaScreen). Two additional reports described twenty-four rare heterozygous variants and performed functional analyses using CRE luciferase reporter assays to generate six-point concentration-response curves (23, 24) (Table 1). However, only seven variants were shown to reduce αMSH-induced MC4R cAMP signaling, while some variants enhanced signaling (23), incongruous with known MC4R inactivation usually observed in obesity.

**Table 1.** Human MRAP2 genetic variants reported in studies of overweight and obesity.

| Variant Type | Variant | <sup>a</sup>Phenotype in report | <sup>b</sup>Functional characterization | <sup>c</sup>Location | Allele count (GnomAD) | Reference |
| --- | --- | --- | --- | --- | --- | --- |
| **Missense** |  |  |  |  |  |  |
|  | **A3T** | 1 ow | WT-like | ECD | 21 | (23) |
|  | **A3S** | 3 ob | WT-like | ECD | 1 | (23) |
|  | **Q13E** | 1 nw | Increased | ECD | 1 | (23) |
|  | **G31V\*** | 1 ob | Decreased | ECD | 0 | (23) |
|  | **P32L\*** | 1 nw | WT-like | ECD | 3 | (23) |
|  | **A40S** | 1 PWL | ND | ECD | 6 | (21) |
|  | **G52R** | 1 ob | Decreased | TM | 6 | (24) |
|  | **F62C\*** | 1 ow | Decreased | TM | 0 | (23) |
|  | **N77S** | 2 ob | Decreased | ICD | 35 | (23) |
|  | **N88Y\*** | 1 ob | ND | ICD | 30 | (15) |
|  | **V91A\*** | 1 nw | Increased | ICD | 5 | (23) |
|  | **E99Q** | 1 ob | WT-like | ICD | 10 | (23) |
|  | **R113G\*** | 2 ob | WT-like | ICD | 15 | (23) |
|  | **S114A\*** | 1 ob | WT-like | ICD |  | (23) |
|  | **L115V\*** | 1 ob | ND | ICD | 70 | (15) |
|  | **N121S\*** | 1 ob | WT-like | ICD | 1 | (23) |
|  | **R125C\*** | 1 ob (15); 1 ob (22); 6 ob, 4 ow, 3 nw (23) | Increased | ICD | 693 | (15, 23) |
|  | **R125H** | 1 ob, 1 nw (22); 15 ob, 10 ow, 5 nw (23); 2 PWL, 2 ob (21) | Increased(23), WT-like(22) | ICD | 1669 | (21-23) |
|  | **H133Y\*** | 1 nw | Increased | ICD | 3 | (23) |
|  | **A137T** | 1 ob, 1 nw (22) 2 nw (23) | Increased(23), WT-like(22) | ICD | 12 | (22, 23) |
|  | **E157G** | 1 ob | ND | ICD | 2 | (29) |
|  | **M162T** | 1 ob, 1 nw | Increased | ICD | 28 | (23) |
|  | **Q174R** | 1 ob | Decreased | ICD | 58 | (22) |
|  | **T193A\*** | 1 ob | Increased | ICD | 2 | (23) |
|  | **P195L** | 2 ob, 1 ow | Decreased | ICD | 4 | (23) |
|  | **D203Y** | 1 nw | Increased | ICD | 23 | (23) |
| **Nonsense** |  |  |  |  |  |  |
|  | **E24X** | 1 ob | ND | N/A | 8 | (15) |
|  | **K102X** | 1 ow | Decreased | N/A | 1 | (23) |

<sup>a</sup>Data was derived from six studies that reported MRAP2 variants in association with overweight (ow), obesity (ob), Prader-Willi-like syndrome (PWL) or normal weight (nw). <sup>b</sup>Two studies (23, 24) assessed MRAP2 effects on MC4R-mediated CRE luciferase responses in CHO cells. MRAP2 and MC4R were expressed in an 8:1 ratio. These studies investigated the effects of αMSH and ACTH. The latter was used based on a zebrafish study that showed MRAP2 can switch MC4R to an ACTH receptor (41). However, as this is a single report, the physiological role of this phenomenon has not been assessed in humans, and to allow comparison between studies, we excluded ACTH data from the table. One study (22) used a cAMP AlphaScreen assay to assess MC4R responses to a single concentration of αMSH (1 μM) in HEK293 cells with MRAP2 and MC4R expressed in a 4:1 ratio. ND, not determined. <sup>c</sup>ECD, extracellular domain, ICD, intracellular domain, TMD, transmembrane domain. Asterisk indicates variants functionally characterized in this study.

There are several reasons why MRAP2 variants may not have affected MC4R signaling in these studies. Firstly, effects may have been missed as previous studies have not used dynamic signaling assays and instead use either single agonist concentrations, end-point assays following cell lysis, or measure parameters downstream of immediate receptor activation (e.g. CRE luciferase reporters that measure transcription). Furthermore, previous studies overexpressed MRAP2 at 4-8x the concentration of MC4R (22–24), and whether MRAP2 variants affect MC4R signaling when expressed at equivalent concentrations is unknown. Additionally, these studies only investigated cAMP signaling, which has been considered the canonical pathway by which MC4R signals. However, ∼25% of MC4R mutations do not affect cAMP signaling, and these have recently been shown to affect G protein coupling, signaling by other pathways (e.g. Gq, ERK1/2), β-arrestin recruitment and internalization (25, 26), and the effect of MRAP2 variants on these other MC4R pathways has not been explored. Clarifying whether MRAP2 variants affect MC4R signaling could have important implications for patient care as the MC4R agonist, setmelanotide, has recently been licensed for use in chronic weight management for genetic obesity syndromes affecting the MC4R pathway (27).

Here we re-examined the effect of twelve previously described MRAP2 variants on MC4R activity. Using dynamic assays, we investigated whether these variants affect MRAP2 protein expression, MC4R cell surface expression and internalization, and MC4R-induced cAMP and IP3 signaling to obtain a more comprehensive understanding of how MRAP2 variants affect MC4R activity.

## Results

### Confirmation that MRAP2 enhances MC4R signaling and impairs trafficking

Previous studies have shown that MRAP2 enhances MC4R-driven signaling when MRAP2 is expressed in excess to the receptor (15). However, our studies of MC3R and MRAP2 (28) and a recent preprint (20) demonstrated that MRAP2 can still enhance signaling when the two proteins are expressed at equal concentrations (20). To confirm this, we transfected HEK293 cells with three different amounts of total DNA, but with equal amounts of HA-HALO-MC4R and either FLAG-MRAP2 or pcDNA (200:200 ng, 50:50 ng, 25:25 ng). This showed that MRAP2 enhanced MC4R-mediated cAMP activity, measured by Glosensor, at the three concentrations (Figure 1A) and that overexpression of MRAP2 is unnecessary to assess its effects on GPCR activity. We also showed that MC4R and MRAP2 colocalise at cell surfaces when transfected at equal DNA concentrations (Figure 1B). All subsequent studies were performed with equal DNA concentrations of MC4R and MRAP2.

**Figure 1.**
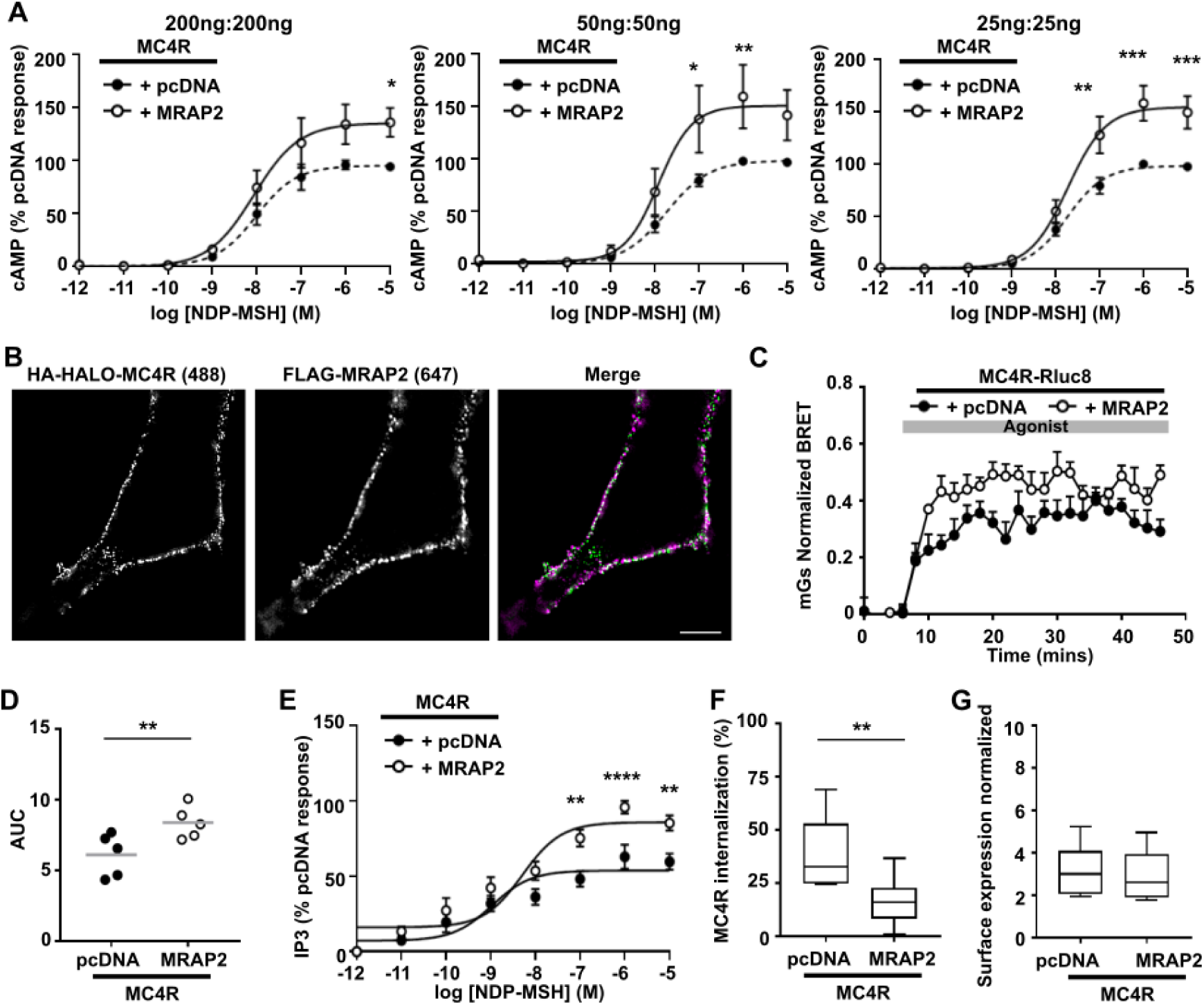
Effect of MRAP2 on MC4R-mediated signaling. (**A**) Measurement of cAMP responses using Glosensor in cells transfected with three different amounts of MC4R with pcDNA or MRAP2. In all assays the same DNA concentration of MC4R and pcDNA or MRAP2 were transfected, shown at the top of the graph. AUC was measured and responses expressed relative to the pcDNA maximal response. N=3. (**B**) SIM showing colocalization between MC4R and MRAP2 at the surface of non-permeabilized HEK293 cells. (**C**) Normalized BRET responses measured between MC4R-Rluc8 and mGs in cells expressing MRAP2 or pcDNA, with (**D**) AUC. N=5. (**E**) MC4R-mediated IP3 responses measured using a NanoBiT biosensor. N=5. (**F**) Percentage internalization of SNAP-MC4R following exposure to NDP-MSH for 30 minutes in cells transfected with pcDNA or MRAP2. N=9. (**G**) Cell surface expression of MC4R assessed by ELISA in cells transfected with HA-SNAP-MC4R and MRAP2 or pcDNA. N=9. Statistical analyses were performed by two-way ANOVA with Sidak’s multiple-comparisons test in A and E, and by student’s unpaired t-test in D, F and G. ****p<0.0001, ***p<0.001, **p<0.01, *p<0.05.

We next sought to confirm recently reported findings that MRAP2 enhances MC4R signaling by both the Gs-cAMP and Gq-IP3 signaling pathways, and that MC4R internalization is impaired by MRAP2 (20). Consistent with the cAMP data, co-expression of MRAP2 with MC4R-Rluc8 enhanced mini-Gs recruitment by BRET when compared to cells co-transfected with pcDNA (Figure 1C-D). Furthermore, MRAP2 significantly enhanced MC4R-driven IP3 responses, assessed using an IP3 NanoBiT Biosensor (Figure 1E). To assess receptor internalization, cells were transfected with HA-SNAP-MC4R, alongside FLAG-MRAP2 or pcDNA, in 96-well plates, exposed to 10 μM NDP-MSH or vehicle for 30 minutes, then surface exposed receptor labeled with membrane impermeable SNAP-surface-647, prior to quantifying the fluorescence. We first showed that the HA-SNAP-MC4R construct trafficked and signaled similarly to the HA-HALO-MC4R (Figure S1A-C). Surface labeling of MC4R was reduced in wells exposed to ligand, consistent with agonist-induced internalization of the receptor and there was significantly greater internalization in cells expressing pcDNA than those expressing MRAP2 (Figure 1F). Finally, we assessed MC4R surface expression using the SNAP-surface-647 in untreated cells. This showed no difference between the surface expression of MC4R in pcDNA or MRAP2 expressing cells (Figure 1G). Thus, these studies confirm that MRAP2 enhances MC4R-driven Gs-cAMP and IP3 responses and impairs receptor internalization.

### Most MRAP2 variants are predicted as pathogenic but do not affect protein expression

Twenty-six missense variants in the MRAP2 gene have been reported in individuals or families with overweight and/or obesity (15, 21–24, 29), although only a subset have been shown to be functionally damaging (Table 1). We selected twelve variants to characterize in detail based on their structural location or variants that present incongruous or inconsistent findings in the literature. We assessed two residues within the extracellular domain (ECD), G31V and P32L, whose functional data has been reported to correlate with their phenotype (loss-of-function in obese individual, WT-like in normal weight individual, respectively) (23), although both are predicted pathogenic (Table 1-2). One highly conserved transmembrane residue, F62C, that reduces cAMP signaling (23) and was identified in an overweight individual (Table 1-2) was also assessed. Nine residues in the intracellular region were selected, seven of which were reported in individuals that were obese or overweight, although all were functionally characterized previously as WT-like or to increase MC4R-induced cAMP signaling (23), inconsistent with MC4R loss-of-function usually observed in obesity (9) (Table 1-2). Most variants had rarely been observed in GnomAD, although the R125C and R125H variants have been observed in 693 and 1669 alleles, respectively (Table 1).

**Table 2.** Pathogenicity prediction for the human MRAP2 variants investigated in this study.

| Human variant | CADD score | PolyPhen-2 | SIFT | Mutation Taster2 | Summary | Evolutionary Conservation<sup>#</sup> |
| --- | --- | --- | --- | --- | --- | --- |
| G31V | 25.3 | Probably damaging | Tolerated | Damaging | 3/4 | 4/6 |
| P32L | 26.9 | Probably damaging | Damaging | Damaging | 4/4 | 6/6 |
| F62C | 28 | Probably damaging | Damaging | Damaging | 4/4 | 6/6 |
| N88Y | 25.5 | Possibly damaging | Damaging | Damaging | 4/4 | 4/6 |
| V91A | 23 | Benign | Tolerated | Damaging | 2/4 | 4/6 |
| R113G | 23 | Probably damaging | Damaging | Damaging | 4/4 | 6/6 |
| S114A | 23.1 | Possibly damaging | Tolerated | Damaging | 3/4 | 6/6 |
| L115V | 16.4 | Benign | Damaging | Damaging | 2/4 | 6/6 |
| N121S | 19.2 | Benign | Damaging | Damaging | 2/4 | 6/6 |
| R125C | 24.8 | Possibly damaging | Damaging | Benign | 3/4 | 2/6 |
| H133Y | 20.1 | Benign | Tolerated | Benign | 1/4 | 4/6 |
| T193A | 19.57 | Benign | Tolerated | Benign | 0/4 | 1/6 |

Protein predictions were made using the online prediction tools CADD (43), Polyphen-2 (44), SIFT (45), MutationTaster2 (46)). The summary column provides a count of the number of tools that predicted the variant to be pathogenic/damaging. CADD integrates several annotations (e.g. conservation, missense changes) into one metric and provides a measure of deleteriousness score, with a high score (>20) representing variants that are not stabilized by selection (i.e. more likely to be disease-causing). PolyPhen-2 integrates sequence and structure-based predictive features to predict damaging mutations (probabilistic score above 0.85 considered probably damaging). SIFT makes predictions about amino acids that will be tolerated based on a combination of homology and amino acid properties. MutationTaster2 predicts a given mutation's likelihood of being disease causing based on SNP frequency and ClinVar data. <sup>#</sup>Evolutionary conservation was assessed between human sequences and mouse, rat, dog, zebrafish-2a, zebrafish-2b, xenopus.

We generated FLAG-tagged constructs of the twelve selected MRAP2 variants and first assessed whether they are predicted to affect the structural conformation of the MRAP2 protein or its expression. No structures of MRAP2 have been published to date, and we therefore generated two models using AlphaFold2, one comprising a single MRAP2 protein, and the second a homodimer, as previous studies have shown both entities to exist at the cell membrane (12, 16, 28, 30). Four of the predicted MRAP2 monomers had additional α-helical structures within the cytoplasmic region, which may lie close to the GPCR intracellular loops and C-terminus, and thus could affect receptor coupling to G proteins or β-arrestin. Several of the ICD residues lie close to or within this cytoplasmic helix and could therefore disrupt G protein or β-arrestin binding, although this is difficult to predict from this model (Figure 2A, Table S2). Within the homodimer structures, the F62 residue is predicted to face into the homodimer interface in two models and forms backbone contacts that are not predicted as affected by mutation to C62 (Figure 2B, Table S2).

**Figure 2.**
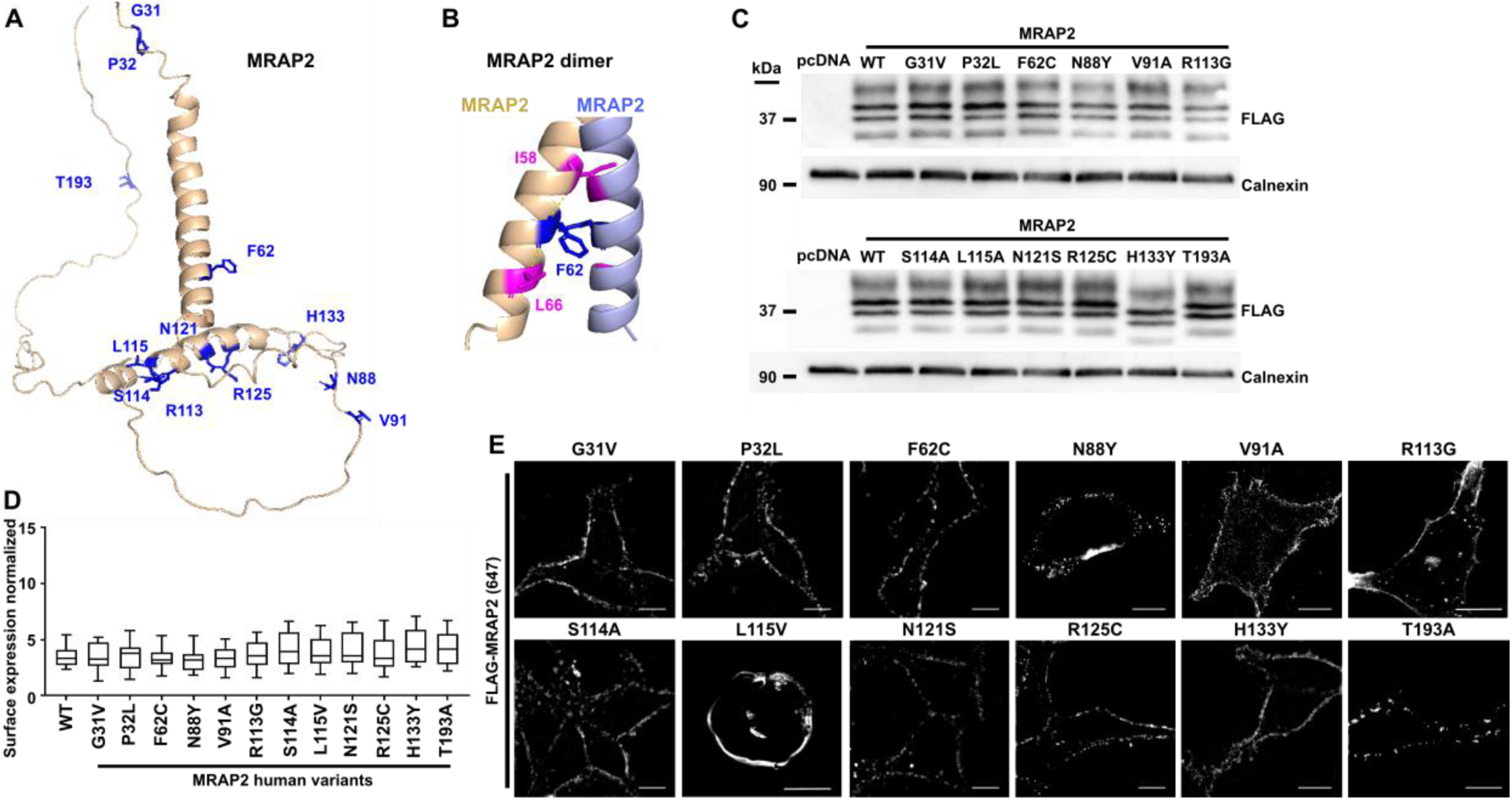
Effect of human variants on MRAP2 protein expression and cell surface expression. (**A**) Structural homology model of the monomeric MRAP2 protein with the locations of the variant residues shown in blue. Two residues are located in the extracellular region, one in the transmembrane region and nine in the intracellular domain. Four of the predicted MRAP2 monomers had α-helical structures in addition to the transmembrane helix where several variant residues are located. These may affect G protein coupling to the receptor. (**B**) Structural model of the homomeric MRAP2 protein showing the F62 residue which forms several backbone contacts. (**C**) Immunoblots showing expression of the FLAG-MRAP2 wild-type and variants in HEK293 cells. Densitometry analysis for N=5-6 blots is shown in Table 3 and full blots in Figure S2. (**D**) Surface expression of FLAG-MRAP2 measured by ELISA in non-permeabilized HEK293. Data is normalized to cells transfected with pcDNA expressed as 0 and not shown. N=10. (**E**) Imaging showing surface expression of MRAP2 variants. Scale, 5 μM.

**Table 3.**
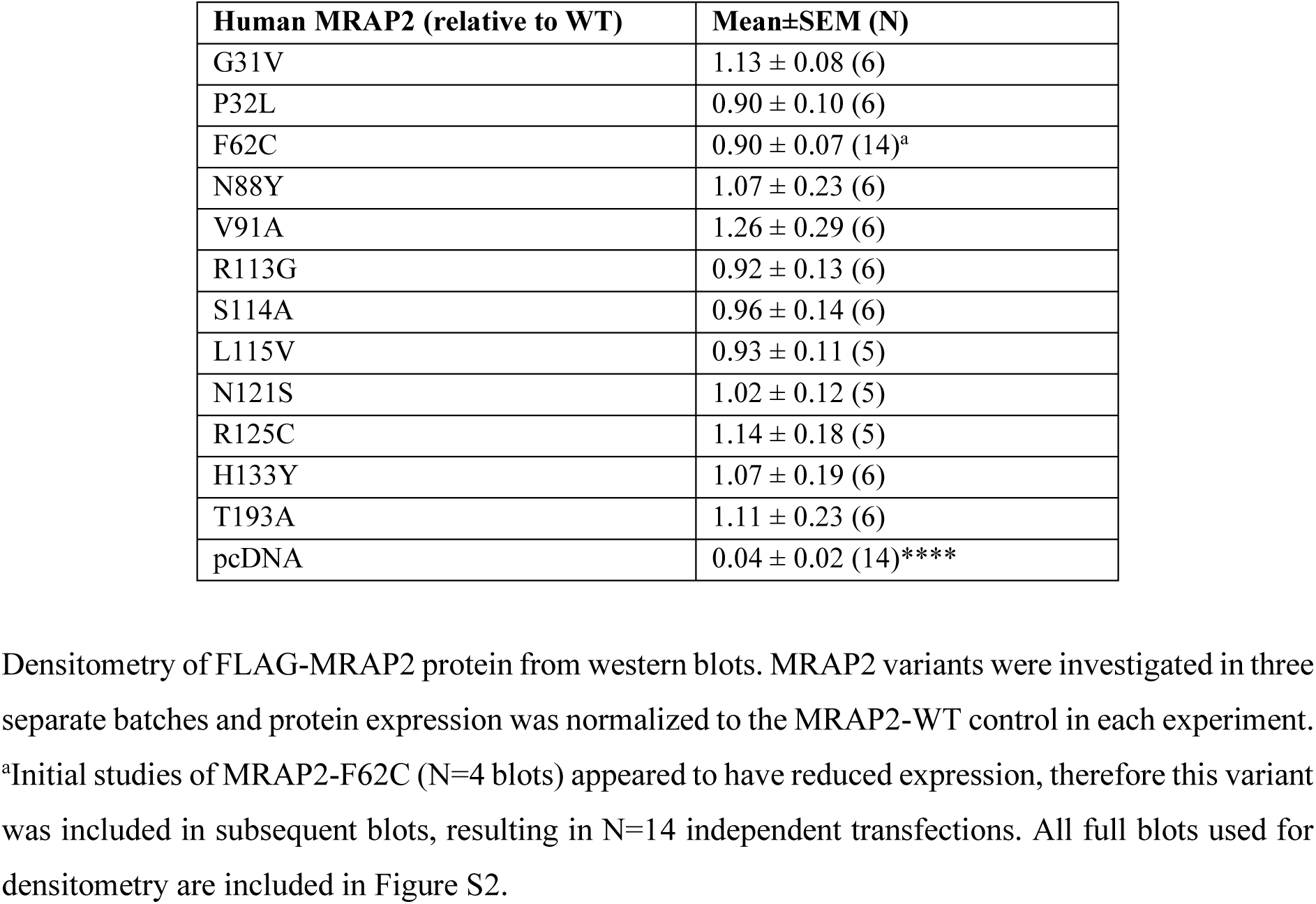
Densitometry of FLAG-MRAP2 protein with human variants.

To determine whether MRAP2 variants affect protein expression we first performed western blot analysis in cells transiently expressing the FLAG-tagged construct. There were no significant differences in the total protein expression (Figure 2C, S2 and Table 3). One variant, H133Y, had a lower apparent molecular weight product than the wild-type (WT) and other MRAP2 variants. However, there was no difference in the isoelectric point and no predicted glycosylation or splice site changes (Table 4), all bands observed in the WT protein were present in the H133Y variant at similar intensities, and whole plasmid sequencing revealed no other differences in plasmid sequence. Although an additional protease cut-site was predicted for MRAP2-H133Y (Table 4), no additional bands at the predicted molecular weight were detected by western blot analysis. Finally, we assessed whether the MRAP2 variants affected MRAP2 cell surface expression in two ways. First, we performed ELISA assays by labeling transfected cells with anti-FLAG antibody and a fluorescent secondary antibody. Quantification of fluorescence on non-permeabilized cells showed no significant difference between the cell surface expression of the variant proteins and WT MRAP2 (Figure 2D). Secondly, the expression of MRAP2 at the surface of non-permeabilized cells was assessed by structured illumination microscopy. This confirmed that all MRAP2 variants were expressed at the cell surface (Figure 2E).

**Table 4.** Predicted effects of the H133Y mutant on MRAP2 isoelectric point, protease sites and glycosylation.

| Variant | Molecular Weight | Isoelectric point | Peptide cutter |  | NetOGlyc | NetNGlyc |
| --- | --- | --- | --- | --- | --- | --- |
|  |  |  | Predicted Proteases | Sites |  |  |
| Wild-type MRAP2 | 26.52 | pH 4.50 | Chymotrypsin, proteinase K, and pepsin | No cleavage site at C132 | 197 (0.716897)<br>199 (0.833337)<br>200 (0.863599)<br>206 (0.653899) | N9 (0.7917) |
| MRAP2 H133Y | 26.55 | pH 4.44 | Chymotrypsin, proteinase K, and pepsin | Predicted cleavage at C132 (detectable fragment: predicted 11.3 kDa) | 197 (0.715397)<br>199 (0.832263)<br>200 (0.862871)<br>206 (0.652319) | N9 (0.7917) |

A single difference was detected between the wild-type MRAP2 and the H133Y mutant MRAP2 with an additional cleavage site predicted resulting in an 11.3kDa fragment of MRAP2-H133Y. However, this fragment was not detected by western blot analyses.

### MRAP2 interacts with TM5-TM6 of the MC4R

We next investigated whether MRAP2 variants are predicted to affect the MC4R-MRAP2 structure. We generated predicted structural models of MRAP2 and MC4R as no published crystal or cryo-EM structures exist of the complex. All five models predicted that MRAP2 interacts with MC4R in a 1-to-1 stoichiometry, consistent with previous studies of MC3R-MRAP2 homodimers (28). MRAP2 was predicted to interact with MC4R TM5 and/or TM6 and identified 8 putative contact sites with the MRAP2 TMD in at least one model (Figure 3A-B Table 5). Alanine mutagenesis of two MRAP2 residues (K42, L64) predicted in at least three models to contact MC4R significantly reduced MC4R-mediated cAMP and IP3 signaling (Figure 3C-D, Table 5). Mutation of the T68 residue had no effect on MC4R signaling (Figure 3C-D). As mutagenesis of predicted contact sites disrupts MC4R signaling, this suggests it is a reliable model to study the effects of MRAP2 variants on MRAP2-MC4R interactions.

**Figure 3.**
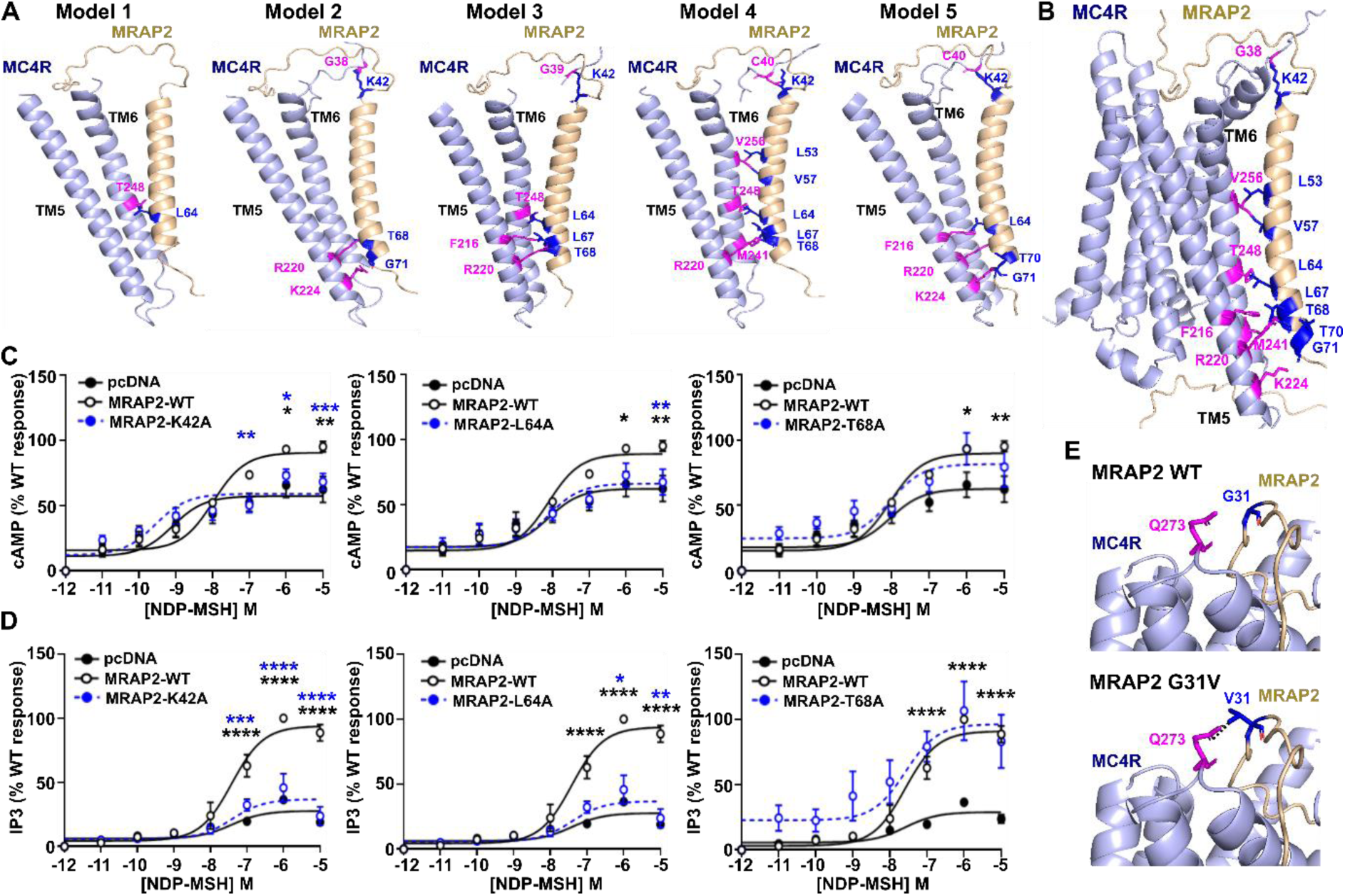
MRAP2 is predicted to interact with MC4R TM5 and TM6. (**A-B**) Predicted structural models of MC4R and MRAP2 in a 1-to-1 configuration with putative interacting residues identified in at least one model highlighted in blue (MRAP2) and pink (MC4R) and detailed in Table 5. (A) Shows the putative interaction sites in all five models between MRAP2 and TM5 and TM6 of MC4R, while (B) shows the full MC4R transmembrane domain structure of model 4. (**C-D**) Effect of alanine mutagenesis of three MRAP2 residues predicted to interact with MC4R on (**C**) cAMP (N=4) and (**D**) IP3 responses (N=3). Statistical analyses show comparisons between pcDNA and WT (black) and WT and alanine mutants (blue). ****p<0.0001, ***p<0.001, **p<0.01, *p<0.05. (**E**) Predicted effects of the MRAP2 G31V mutant on the MC4R-MRAP2 structural model. The mutant V31 residue is predicted to gain a contact with a neighboring residue in MC4R.

**Table 5.** Identification of possible contacts between the MRAP2 transmembrane helix and MC4R in AlphaFold2 models.

| MRAP2 | MC4R |  |  |  |  | No. of models<sup>a</sup> |
| --- | --- | --- | --- | --- | --- | --- |
|  | Model 1 | Model 2 | Model 3 | Model 4 | Model 5 |  |
| K42* |  | G38 (ECD) | G39 (ECD) | C40 (ECD) | C40 (ECD) | 4 |
| L53 |  |  |  | V256 (TM6) |  | 1 |
| V57 |  |  |  | V256 (TM6) |  | 1 |
| L64* | T248 (TM6) |  | T248 (TM6) | T248 (TM6) | F216 (TM5),<br>R220 (TM5) | 4 |
| L67 |  |  | F216 (TM5),<br>R220 (TM5) | R220 (TM5) |  | 2 |
| T68* |  | R220 (TM5) | R220 (TM5) | M241 (TM6) |  | 3 |
| T70 |  |  |  |  | K224 (TM5) | 1 |
| G71 |  | K224 (TM5) |  |  | K224 (TM5) | 2 |

We then examined the effects of the MRAP2 variants on the MRAP2:MC4R structural model. Most variants are located in unstructured regions of the MRAP2, therefore their effects on the MC4R structure cannot be predicted. Three residues are in close proximity to MC4R. G31 and P32 are at the top of the MC4R structure within a cleft above the ligand-binding site (31). The G31V variant is predicted to gain contacts with a neighbouring residue, Q273, in an extracellular loop of MC4R, which could reduce the receptor’s flexibility or impair access to the ligand-binding region (Figure 3E). The P32L variant is predicted to have no effect on the MRAP2-MC4R structure. As F62 faces away from the MC4R TMD, it is difficult to predict the effect of mutation of this residue. However, it is possible that the change from a non-polar phenylalanine to a polar cysteine at residue 62 could affect the residues behaviour, or interactions with the surrounding phospholipid bilayer.

### Some MRAP2 variants impair MC4R internalization

Whether MRAP2 variants affect MC4R expression has not been examined. To determine whether MRAP2 affects MC4R cell surface expression cells were transfected with equal concentrations of MRAP2-WT or the variant plasmids and HA-SNAP-MC4R, then surface exposed receptors labeled with SNAP-surface-647. Fluorescence quantification revealed no significant difference between the cell surface expression of MC4R on cells expressing either MRAP2-WT, MRAP2 human variants or pcDNA (Figure 4A).

**Figure 4.**
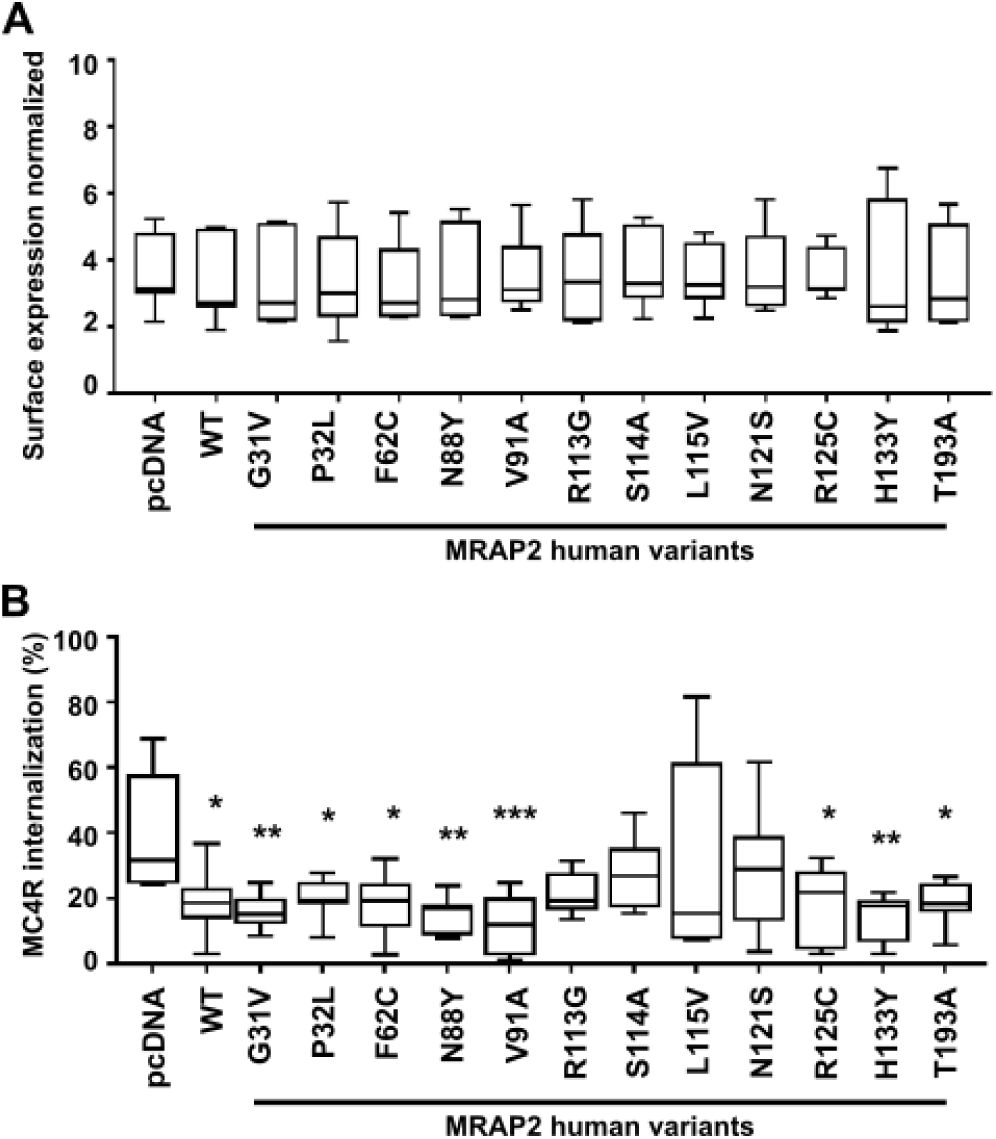
Effect of MRAP2 human variants on MC4R cell surface expression and internalization. (**A**) Surface expression of FLAG-MC4R expressed alongside MRAP2 human variants, measured by ELISA in non-permeabilized HEK293. N=9. (**B**) MC4R-induced internalization in cells expressing pcDNA, MRAP2 wild-type or the twelve MRAP2 variants following 30-minute exposure to 10 μM NDP-MSH. N=8. Statistical analyses were performed by one-way ANOVA with Dunnett’s multiple-comparisons test. Asterisks show comparisons to pcDNA in black. ***p<0.001, **p<0.01, *p<0.05.

As our studies had shown that MRAP2 reduces MC4R internalization, we hypothesized that some MRAP2 variants may impair this effect. To assess receptor internalization, cell surface receptor was quantified following exposure to vehicle or 10μM NDP-MSH in cells expressing HA-SNAP-MC4R, alongside pcDNA, FLAG-MRAP2-WT or MRAP2 variants. Eight variants had similar levels of internalization to wild-type MRAP2. However, MRAP2-R113G, -S114A, -L115V and -N121S had increased internalization such that they were not significantly different to cells expressing no MRAP2 (pcDNA control) (Figure 4B, Table 6). Therefore, MRAP2 variants in the cytoplasmic region are more likely to impair MRAP2s ability to reduce MC4R internalization.

**Table 6.** Summary of the effect of MRAP2 human variants on MC4R signaling compared to wild-type MRAP2.

| **Variant** | **Surface expression** | **Internalization** | **cAMP** | **IP3** |
| --- | --- | --- | --- | --- |
| **G31V** | WT-like | WT-like | Decrease | Decrease |
| **P32L** | WT-like | WT-like | WT-like | WT-like |
| **F62C** | WT-like | WT-like | Decrease | Decrease |
| **N88Y** | WT-like | WT-like | Decrease | Decrease |
| **V91A** | WT-like | WT-like | WT-like | WT-like |
| **R113G** | WT-like | Increase | Decrease | Decrease |
| **S114A** | WT-like | Increase | Decrease | Decrease |
| **L115V** | WT-like | Increase | Decrease | Decrease |
| **N121S** | WT-like | Increase | Decrease | Decrease |
| **R125C** | WT-like | WT-like | WT-like | Decrease |
| **H133Y** | WT-like | WT-like | WT-like | WT-like |
| **T193A** | WT-like | WT-like | WT-like | Decrease |

### MRAP2 variants impair MC4R-mediated cAMP and IP3 signaling

MC4R has been described to predominantly signal by the Gs-cAMP pathway, and ∼75% mutations in MC4R that are associated with severe early onset obesity impair cAMP signaling (25). In contrast, previous studies of MRAP2 variants have shown very few impair MC4R cAMP signaling, although dynamic cAMP responses in live cells were not assessed (22–24). To determine whether MRAP2 variants affect MC4R canonical signaling, cAMP Glosensor assays were performed that are capable of quantifying agonist-induced cAMP responses in real-time in live cells. Seven MRAP2 variants (G31V, F62C, N88Y, R113G, S114A, L115V, N121S) significantly reduced MC4R-mediated cAMP signaling when compared to cells expressing MRAP2-WT. The five other variants had no effect on cAMP signaling (i.e. they were WT-like) (Figure 5, Table 6). Four of these WT-like variants have previously been described in individuals with normal weight (Table 1) and therefore may not have a role in the pathogenesis of weight gain.

**Figure 5.**
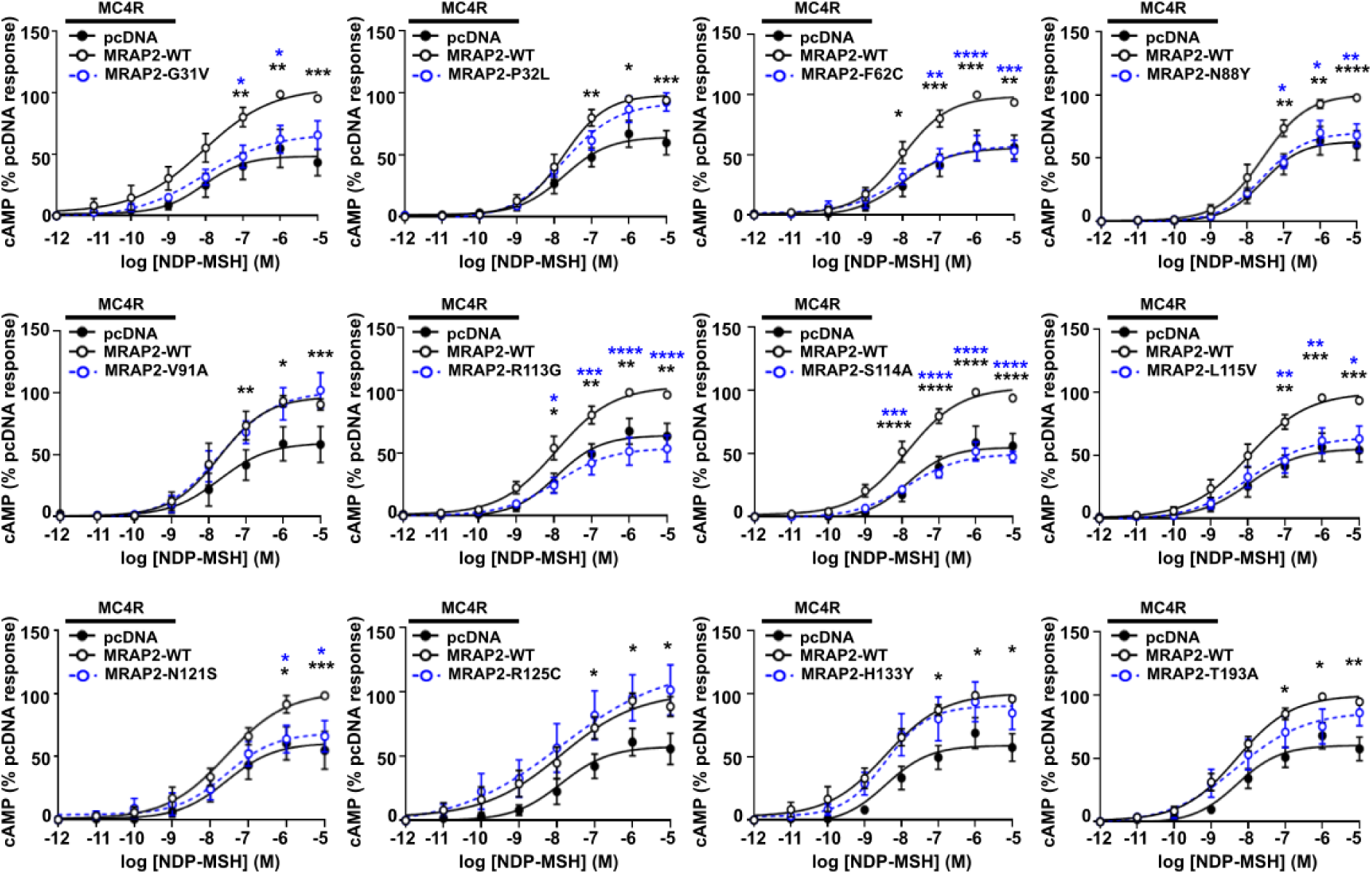
Effect of MRAP2 human variants on MC4R-mediated cAMP signaling. MC4R-mediated cAMP responses in cells transfected with pcDNA, MRAP2-WT or: MRAP2-G31V (N=6), MRAP2-P32L (N=7), MRAP2-F62C (N=7), MRAP2-N88Y (N=6), MRAP2-V91A (N=4), MRAP2-R113G (N=7), MRAP2-S114A (N=8), MRAP2-L115V (N=8), MRAP2-N121S (N=5), MRAP2-R125C (N=6), MRAP2-H133Y (N=6), MRAP2-T193A (N=7).). Statistical analyses show comparisons between pcDNA and WT (black) and WT and alanine mutants (blue) performed by two-way ANOVA with Sidak’s multiple-comparisons test. ****p<0.0001, ***p<0.001, **p<0.01, *p<0.05.

A subset of obesity-associated MC4R variants impair Gq-IP3 signaling in addition to, or instead of, Gs-cAMP signaling (25). To determine whether MRAP2 variants affect MC4R-mediated IP3 signaling, NDP-MSH-induced responses were measured using a NanoBiT IP3 biosensor in live cells. Seven MRAP2 variants (G31V, F62C, N88Y, R113G, S114A, L115V, N121S) that impaired MC4R-induced cAMP signaling also reduced MC4R-mediated IP3 responses (Figure 6, Table 6). Two additional MRAP2 variants (R125C and T193A) also reduced MC4R-mediated IP3 signaling, while the other three variants had no effect on IP3 signaling (Figure 6). Therefore, MRAP2 variants can affect multiple aspects of MC4R signaling (Table 6).

**Figure 6.**
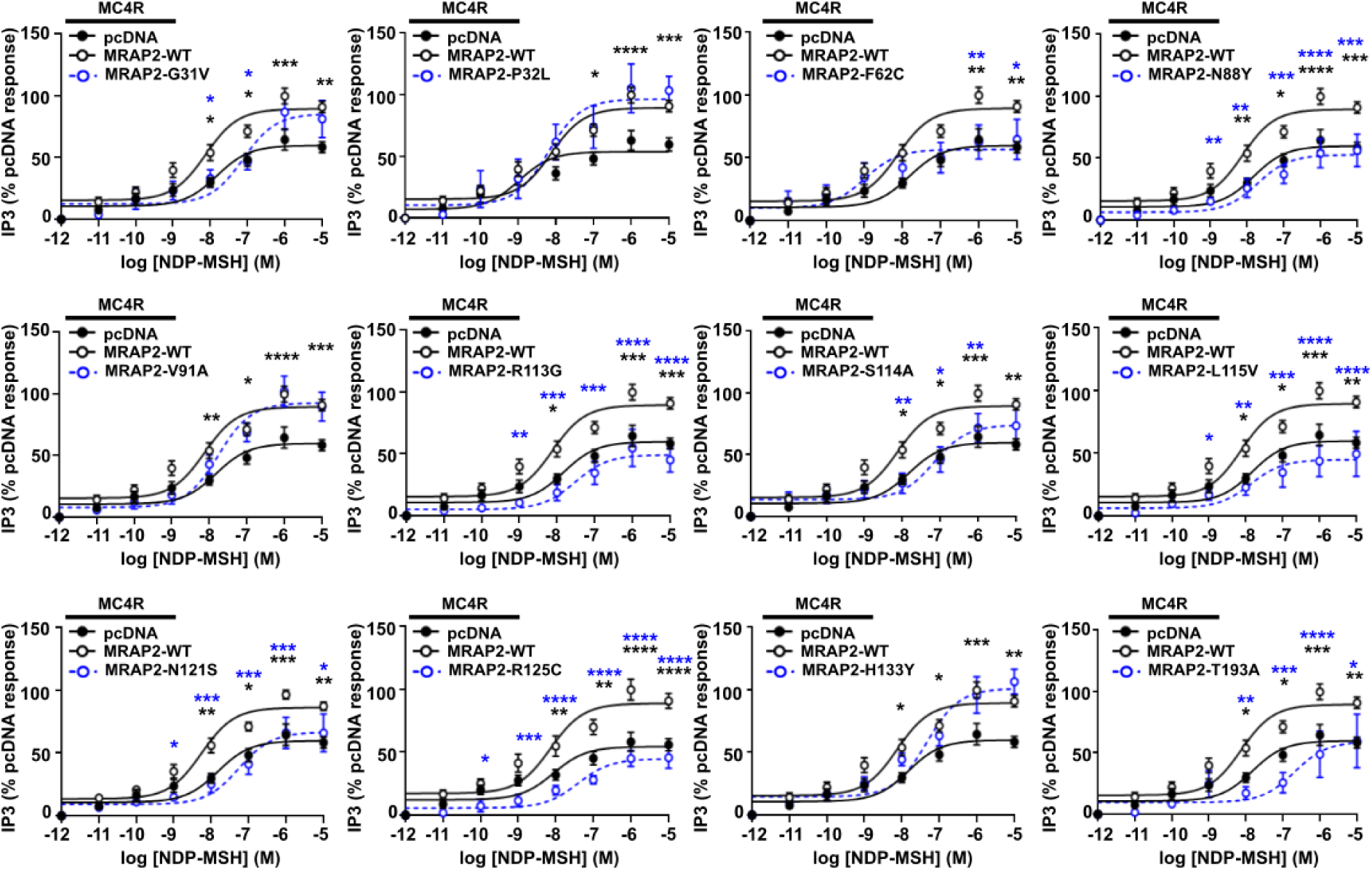
Effect of MRAP2 human variants on MC4R-induced IP3 signaling. MC4R-mediated IP3 responses in cells transfected with pcDNA, MRAP2-WT or: MRAP2-G31V (N=5), MRAP2-P32L (N=5), MRAP2-F62C (N=5), MRAP2-N88Y (N=5), MRAP2-V91A (N=5), MRAP2-R113G (N=4), MRAP2-S114A (N=5), MRAP2-L115V (N=5), MRAP2-N121S (N=6), MRAP2-R125C (N=4), MRAP2-H133Y (N=5), MRAP2-T193A (N=5). Statistical analyses show comparisons between pcDNA and WT (black) and WT and alanine mutants (blue) performed by two-way ANOVA with Sidak’s multiple-comparisons test. ****p<0.0001, ***p<0.001, **p<0.01, *p<0.05.

## Discussion

Our studies have demonstrated that MRAP2 variants associated with human obesity impair multiple aspects of MC4R signaling. Consistent with previous studies we showed that wild-type MRAP2 enhances MC4R coupling to Gs and increases cAMP signaling (15, 20), and demonstrated that most obesity-associated MRAP2 variants impair cAMP signaling. However, some variants did not affect cAMP signaling which is consistent with previous studies of obesity-associated MC4R mutations (25, 26). All MRAP2 variants that were identified in at least one overweight or obese individual also impaired MC4R-mediated IP3 signaling. Although the cAMP pathway has been considered the canonical MC4R signaling pathway for many years, studies in the last decade have demonstrated the importance of Gq/11 signaling for MC4R physiological responses. Thus, mice lacking Gq/11 at the paraventricular nucleus of the hypothalamus develop severe hyperphagic obesity and are unable to respond to MC4R agonists (32), while obese mice with a MC4R-F51L mutation associated with human obesity, have a specific defect in Gq/11 signaling (33). Moreover, impairment of other signaling pathways including Gq and ERK1/2 pathways have been described for several obesity-associated MC4R mutations (25, 26) in cellular studies. In combination, these studies suggest that the Gq/11-IP3 signaling pathway may be as important as the Gs-cAMP pathway, and both should be studied to understand the impact of variants in MC4R and MRAP2.

Previous studies of some of the MRAP2 variants that we investigated did not detect any differences in MC4R signaling or enhanced signaling, incongruous with MC4Rs known role in weight regulation (23). There were several differences between our studies which could explain this variability. Firstly, we performed all our assays in the presence of equal DNA concentrations of MC4R and MRAP2, whereas the previous studies overexpressed MRAP2 at a concentration 4-8 times higher than MC4R. We used equal DNA concentrations as our studies showed the accessory protein could still enhance signaling under these conditions, as did a recent preprint (20, 22), and we could not identify any rationale in the literature for overexpression of MRAP2. Future studies of the effects of MRAP2 on GPCRs should similarly avoid significant overexpression. Our studies also performed different signaling assays to detect changes in MC4R signaling compared to the previous studies which either did not perform full concentration-response curves, used endpoint assays far downstream of receptor activation or only investigated the cAMP pathway as this has previously been considered to be the canonical MC4R pathway (22, 23). Therefore, it will be important to capture dynamic signaling behaviour with a range of agonist concentrations to determine the effects of MRAP2 variants on GPCR behaviour in future studies.

One MRAP2 variant, H133Y, consistently produced a smaller protein product on immunoblot analyses. However, no differences in the predicted isoelectric point, glycosylation or splice sites were predicted between the mutant and wild-type proteins. Although an additional protease site was predicted for the MRAP2-H133Y protein, we did not detect smaller products of this size by western blot analyses. Functional characterization of the effect of the H133Y mutant on MC4R and MC3R signaling and trafficking (28) detected no differences when compared to MRAP2 wild-type proteins. As this mutant protein has only been detected in normal weight individuals, it is likely to be a benign single nucleotide variant that has no functional effect on MRAP2. Two other MRAP2 variants, P32L and V91A, also had no effect on MC4R activity in our studies and were previously shown to either enhance or not affect MC4R activity (23). As these variants also had no effect on MC3R signaling in our previous studies (28) and have only been reported in normal weight individuals they are unlikely to contribute significantly to obesity pathogenesis. The R125C MRAP2 mutant has also been identified in overweight and obese individuals as well as normal weight family members. We showed that it reduces MC4R-mediated IP3 responses and a previous study has suggested mutation of the R125 residue affects MRAP2 glycosylation (34). Therefore, this mutation likely has reduced penetrance, which has also been observed for some MC4R mutations (9).

Four MRAP2 mutations enhanced MC4R internalization when compared to wild-type MRAP2, and disrupted both cAMP and IP3 signaling. These mutant residues are located intracellularly and are predicted to form part of a helical structure that lies perpendicular to the transmembrane domains of MRAP2 and MC4R. This is analogous to our previous study of MC3R and MRAP2 complexes in which we identified a similar intracellular helical structure that lies close to the intracellular loops and mutations in the four MRAP2 residues similarly affected internalization and signaling of MC3R (28). The juxtamembrane region where this helical structure is located is important for the conformational rearrangements required for MC4R-G protein coupling, which involves interactions with TM3, TM5-7 and all three intracellular loops (35, 36). MC4R mutations within these intracellular loops have been shown to disrupt receptor internalization and β-arrestin recruitment (25, 37). It is likely that residues within the MRAP2 α-helix facilitate G protein coupling and that mutations in this structure disrupt these processes resulting in impaired receptor signaling.

Our studies predicted that MRAP2 interacts with residues in MC4R TM5 and TM6. A preprint has also identified similar interactions between MRAP2 and MC4R TM5 or TM6 in a homology model based on the MC2R-MRAP1 structure (20). Moreover, our previous studies of MC3R and MRAP2 identified several of the same MRAP2 residues (K42, L53, L64, L67 and T68) to form contacts with TM5 and/or TM6 of MC3R (28). Therefore, this suggests that MRAP2 may interact with melanocortin receptors using a shared mechanism. As previous studies have shown that MC4R activation requires outward displacement of the intracellular end of TM6 and inward movement of TM5 to allow G protein coupling (35, 36, 38), we predict that MRAP2 binding to melanocortin receptors must facilitate a GPCR conformation that allows this active conformation to be adopted more readily than receptor in the absence of the accessory protein. Additionally, our structural models suggest that MRAP2 also interacts with residues in the extracellular N-terminal region of MC4R. These interactions involve the MRAP2-K42 residue and we showed that mutation of this residue impairs MC4R-mediated Gs-cAMP and Gq-IP3 signaling. MRAP2-K42 is predicted to interact with MC4R-C40, a residue that forms an intra-disulfide bond with C279 in extracellular loop-3 in active MC4R-Gs-bound structures, but not in inactive states (36). Mutations in C40 have been reported in individuals with obesity (39, 40), and we predict that MRAP2 interactions with C40 could be important in maintaining the structural integrity of the MRAP2-MC4R-Gs bound state.

MRAP2 had no effect on the overall cell surface expression of MC4R in our studies, and we did not detect any differences in MC4R expression in the presence of MRAP2 mutant proteins. This is consistent with previous studies that similarly reported that MRAP2 does not affect MC4R surface expression (20, 41). However, some studies have suggested MRAP2 can reduce (12, 16) or increase MC4R cell surface expression (18), while investigations in inner medullary collecting duct (IMCD3) cells have shown that MRAP2 enhances MC4R expression at primary cilia, an organelle that is known to be important in energy homeostasis and weight control (19). As we did not examine ciliated cells we cannot exclude the possibility that MRAP2 mutant proteins may alter expression of MC4R at primary cilia and this would be important to characterise in future studies. Despite detecting no effects of MRAP2 on the overall MC4R cell surface expression, we did show that MRAP2 impairs receptor internalization (Figure 1), which could be one mechanism by which MRAP2 enhances receptor signaling. Moreover, we showed that obesity-associated MRAP2 mutations impair this effect, and consequently have increased MC4R internalization, which could explain the reductions in MC4R signaling by these MRAP2 mutant proteins. However, MRAP2 mutants that impair MC4R signaling do not always enhance receptor internalization, and therefore other mechanisms must exist to explain the effects of these MRAP2 mutant proteins on receptor activity.

In summary, our studies demonstrate that human MRAP2 variants that are associated with obesity impair MC4R signaling by affecting multiple signaling pathways. Future studies of MRAP2 variants should assess their effects on both Gs-cAMP and Gq-IP3 pathways to determine their pathogenicity.

## Materials & Methods

### Plasmid constructs and compounds

A list of plasmids with their source can be found in Table S1. MC4R constructs were generated within a pRK5 vector with an mGluR2 N-terminal signal peptide (42), followed by affinity tags (HA or FLAG), self-labeling protein tags capable of conjugation to organic dyes (SNAP or Halo), followed by human MC4R. Cloning into the pRK5 vector was performed using reagents from Promega and oligonucleotides from Sigma to generate ss-HA-Halo-MC4R and ss-HA-SNAP-MC4R. All plasmids were sequence-verified by Source Bioscience. NDP-MSH (Cambridge Bioscience) was used at a concentration of 10 µM, unless otherwise stated. Whole plasmid sequencing was performed using Oxford Nanopore sequencing at plasmidsNG (The BioHub, Birmingham).

### Cell culture and transfection

Adherent HEK293 (Agilent Technologies) were maintained in DMEM-Glutamax media (Merck) with 10% calf serum (Invitrogen) at 37 °C, 5% CO_2_. Cells were routinely screened to ensure they were mycoplasma-free using the TransDetect Luciferase Mycoplasma Detection kit (Generon). Expression plasmids were transiently transfected into cells using Lipofectamine 2000 (Life Technologies).

### Protein sequence alignment, pathogenicity predictions and three-dimensional modeling of MC4R and MRAP2

CADD (43), Polyphen-2 (44), SIFT (45) and MutationTaster2 (46) were used to predict the effect of amino acid substitutions (45–48). Amino acid conservation was examined in MRAP2 mammalian orthologs using ClustalOmega (49). Modeling of MRAP2 monomers, dimers and heterodimers with MC4R was performed using AlphaFold2 with the ColabFold v1.5.2-patch in Google Co-laboratory (50) and visualized using Pymol. FASTA sequences were obtained from NCBI. Five models were predicted and ranked based on predicted local distance difference test (pLDDT). Figures were prepared using the PyMOL Molecular Graphics System (Version 2.5.2, Schrodinger, LLC). H133Y mutant was additionally analysed using Protein Isoelectric Point and Molecular Weight calculators (Sequence Manipulation Suite) (51), Alternative Splice Site Predictor (52), Expasy PeptideCutter (53), and glycosylation was predicted using NetOGlyc-4.0.0.13 and NetNGlyc-1.0 (54) (accessed 10/09/24).

### Assessment of cell surface expression and internalization

AdHEK cells were seeded at 10,000 cells/well in black 96-well plates and transfected the same day with 100 ng pcDNA or 100 ng MRAP2 (WT or variants) only and with 100ng HA-SNAP-MC4R for receptor studies. For cell surface expression, cells were fixed 48-hours later in 4% PFA (Fisher Scientific UK) in PBS, then labeled with 1:1000 anti-FLAG mouse monoclonal antibody (M2, Sigma-Aldrich Cat# F3165, RRID:AB_259529), followed by Alexa Fluor 647 anti-mouse secondary antibody (Cell Signaling Technology Cat# 4410, RRID:AB_1904023). Cells were washed, then fluorescence read on a Glomax plate reader (Promega). Internalization assays were performed 48-hours post-transfection and cells were exposed to 10 μM NDP-MSH or vehicle for 30 minutes, then labeled with SNAP-surface-647.

### Bioluminescence resonance energy transfer (BRET)

NanoBRET assays were performed using methods that have previously been described in detail (55). AdHEK cells were seeded at 100,000 cells/well in 6-well plates and transfected 24-hours later with 100 ng each of MC4R-Rluc8, Venus-mGs and pcDNA or FLAG-MRAP2. Forty-eight hours later, cells were reseeded in 96-well plates in FluoroBrite DMEM phenol red-free media with 10% FBS and 2 mM L-glutamine (FluoroBrite complete media) and incubated at 37 °C, 5% CO_2_ for at least four hours. Media was then replaced with coelenterazine-H (Promega) added at a 1:100 dilution and vehicle and agonists were prepared in HBSS at 10x concentration. BRET measurements were recorded using a Promega GloMax microplate reader at donor wavelength 475-30 and acceptor wavelength 535-30 at 37 °C and the BRET ratio (acceptor/donor) was calculated for each time point. Four baseline recordings were made, then agonist added at 8 minutes and recordings made for a further ∼40 minutes. The average baseline value recorded prior to agonist stimulation was subtracted from the experimental BRET signal. All responses were then normalized to that treated with vehicle to obtain the normalized BRET ratio. AUC was calculated in GraphPad Prism

### Glosensor cAMP assays

AdHEK cells were plated in 6-well plates and transfected 24-hours later with 200 ng pGloSensor-20F plasmid, and equal amounts of MC4R and MRAP2 (25-200 ng for transfection tests, and 25 ng for all other studies). Forty-eight hours later, cells were seeded in 96-well plates in FluoroBrite complete media (ThermoScientific). Cells were incubated for at least 4 hours, then media changed to 50 µL of equilibration media consisting of Ca^2+^- and Mg^2+^-free HBSS containing a 2% (v/v) dilution of the GloSensor cAMP Reagent stock solution. Cells were incubated for 2 hours at 37°C. Basal luminescence was read on a Glomax plate reader for 8 minutes, then agonist added and plates read for a further 30 minutes. Data was plotted in GraphPad Prism, area-within the curve calculated and these values used to plot concentration-response curves with a 4-parameter sigmoidal fit.

### IP3 assays

Assays were performed as previously described (56). AdHEK cells were plated in white 96-well plates and transfected with 200 ng LgBiT-IP_3_R2-SmBiT plasmid, 25 ng HA-Halo-MC4R and either 25ng pcDNA3.1 or 3xFLAG-MRAP2. Following 48 hours, media was changed to 50μl Hank’s buffered saline solution (HBSS), before loading each well with 40 µL NanoGlo substrate (1:100 dilution, Promega) and luminescence read on a Glomax plate reader at 37°C. Vehicle and agonists were prepared in HBSS at 10x concentration and added to wells following recording of baseline signals for four cycles (equivalent to 8 minutes), then responses read for a further 22 minutes. The average baseline value recorded prior to agonist stimulation was subtracted from the experimental signal, then data normalized to vehicle-treated cells. AUC was calculated in GraphPad Prism and these values used to plot concentration-response curves with a 4-parameter sigmoidal fit.

### Western blot analysis

Cells were seeded into 6-well plates then transfected 24-hours later with either 3xFLAG-MRAP2-WT or 3xFLAG-MRAP2-mutants at 1µg per well. Cells were lysed after 48-hours in NP40 buffer and western blot analysis performed, as described (57). Blots were blocked in 5% marvel/TBS-T, then probed with anti-FLAG (M2, Sigma-Aldrich Cat# F3165, RRID:AB_259529) and anti-calnexin (Millipore Cat# AB2301, RRID:AB_10948000) antibodies, the latter after visualizing and stripping the blot following FLAG probing. Blots were visualized using the Immuno-Star WesternC kit (BioRad) on a BioRad Chemidoc XRS+ system. Densitometry was performed using ImageJ (NIH), and protein quantities normalized to calnexin.

### Structured illuminated microscopy (SIM)

Cells were plated on 24 mm coverslips (VWR) and transfected with 500 ng plasmid. For expression of MRAP2, cells were fixed with 4% PFA in PBS and exposed to 1:1000 anti-FLAG mouse monoclonal antibody (M2, Sigma), followed by Alexa Fluor 647 anti-mouse antibody. For internalization studies, cells were transfected with HA-SNAP-MC4R or HA-Halo-MC4R and pcDNA or FLAG-MRAP2 plasmid for 48 hours. Cells were either fixed in 4% PFA in PBS (time point 0) or exposed to 10μM NDP-MSH for 30 minutes before fixation. All coverslips were then exposed to 1:1000 anti-HA mouse monoclonal antibody (BioLegend Cat#901514, RRID:AB_2565336), then 1:3000 Alexa Fluor 647 secondary antibody. Samples were imaged on a Nikon N-SIM system (Ti-2 stand, Cairn TwinCam with 2 × Hamamatsu Flash 4 sCMOS cameras, Nikon laser bed 488 and 647 nm excitation lasers), Nikon 100 × 1.49 NA TIRF Apo oil objective. SIM data was reconstructed using NIS-Elements (v. 5.21.03) slice reconstruction.

### Statistical analysis

Statistical tests used for each experiment are indicated in the legends of each figure and the number of independent biological replicates denoted by N. Data was plotted and statistical analyses performed in Graphpad Prism 7. Normality tests (Shapiro-Wilk or D’Agostino-Pearson) were performed on all datasets to determine whether parametric or non-parametric statistical tests were appropriate. A p value of <0.05 was considered statistically significant.

## Author contributions

Conceptualization: CMG

Methodology: CMG

Investigation: RAW, AJ, VM, CQ, CMG

Writing – original draft: CMG

Writing – review and editing: All authors

## Data and materials availability

All data needed to evaluate the conclusions in the paper are present in the paper and/or the Supplementary Materials. Plasmid constructs developed for this manuscript will readily be made available upon request. Plasmid constructs obtained from other researchers are detailed in Table S1 and may be subject to Material Transfer Agreements. Please contact the corresponding author of this manuscript, or the named source for details.

