## Supplementary Figures for "Obesity-associated MRAP2 variants impair multiple MC4R-mediated signaling pathways"

**Supplementary Appendix**

**Figure S1 MC4R plasmids express, traffic and signal normally**

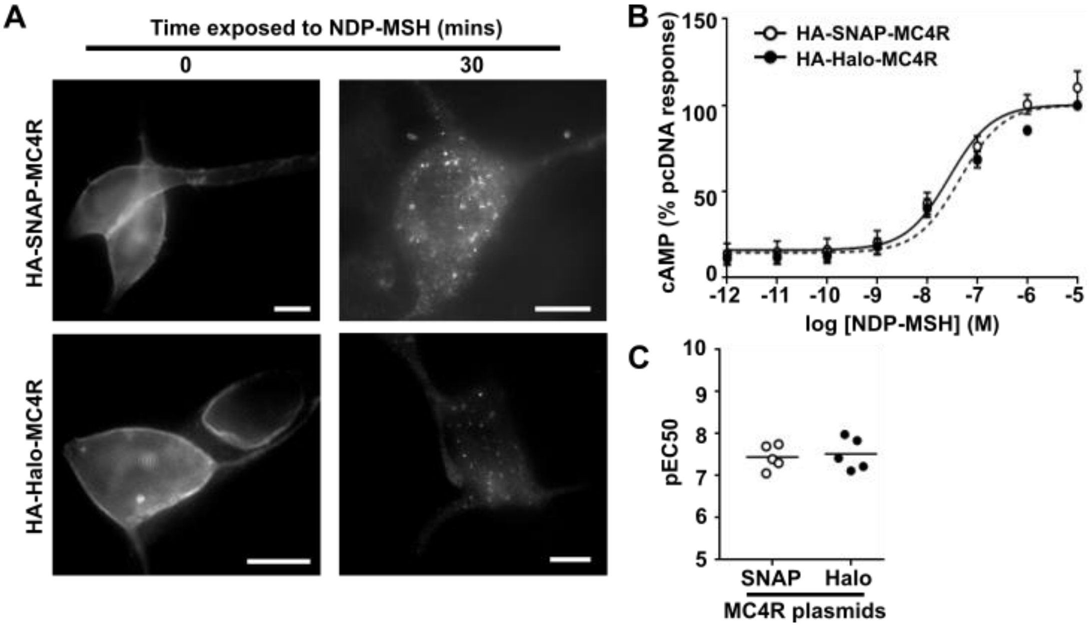


(**A**) Imaging showing cell surface expression and agonist (10 μM NDP-MSH) induced trafficking of the two MC4R constructs. Scale, 5 μm. Cell surface labelling is reduced and more vesicles are present after exposure to agonist for 30 minutes. (**C**) MC4R-induced cAMP responses measured by Glosensor in cells transfected with the two MC4R plasmids and (**D**) pEC50. AUC was used to generate a dose-response and expressed relative to basal responses. N=5. The plasmids traffic normally and elicit similar signaling responses.

**Figure S2 Full blots used for densitometry analysis**

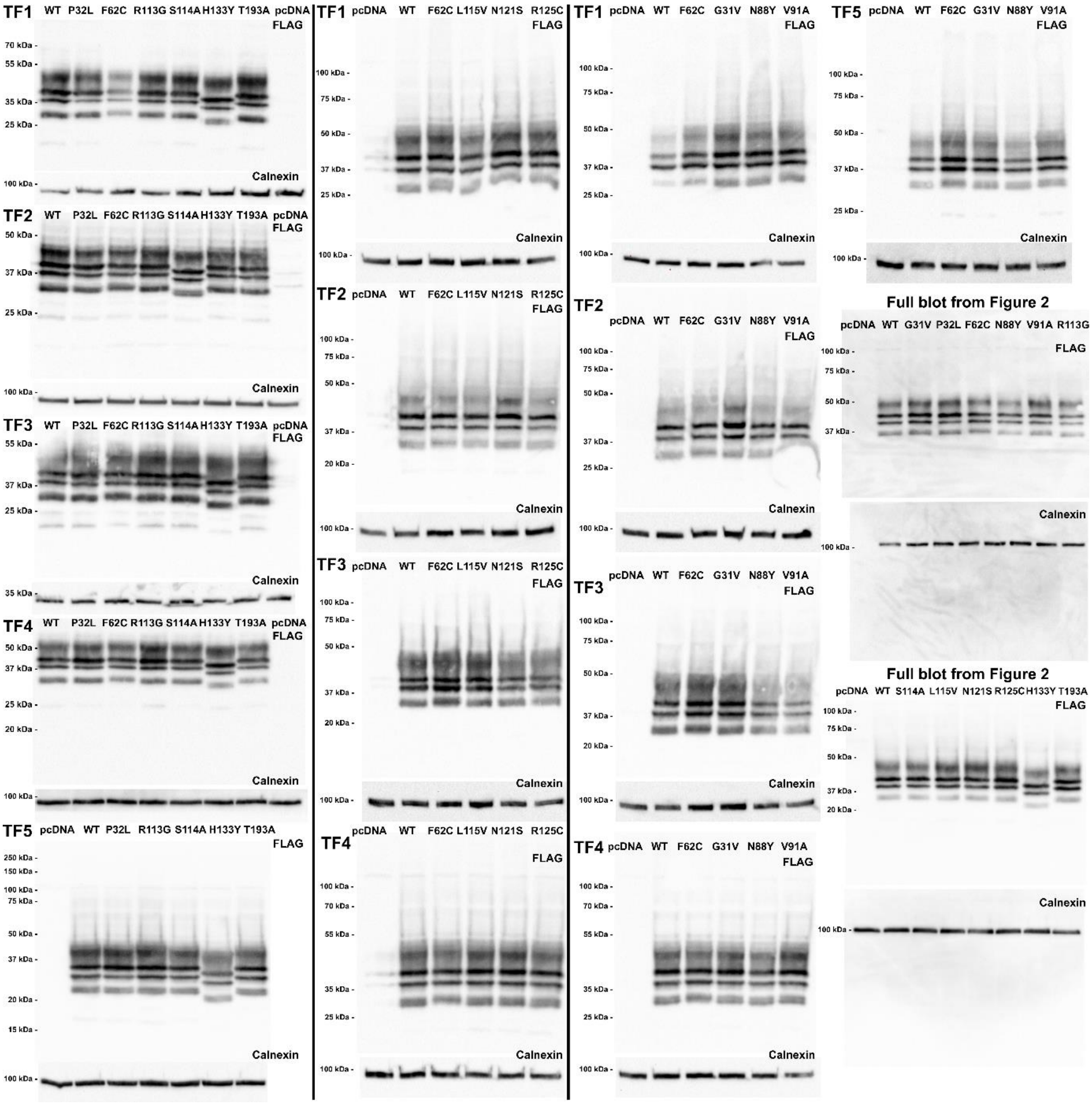


Western blot analyses were performed in three batches shown in columns separated by black lines. Additionally, the full blots from Figure 2 are shown. MRAP2-F62C was included in the first batch of blots (left-hand column) and appeared to have lower expression and was therefore included in subsequent analyses. Blots for transfection (TF) 2, 3 and 4 in the left-hand column were cut before processing and the top part used for the loading control (calnexin) and the bottom used for FLAG-MRAP2 detection.

### Table S1 Expression plasmids used in this manuscript

| **Plasmid name** | | **Information** | | **Source** |
|---|---|---|---|---|
| ss-HA-Halo-MC4R | | N-terminal signal peptide from mGluR5, followed by HA, HALO and human MC4R | | Caroline Gorvin, University of Birmingham (1) |
| ss-HA-SNAP-MC4R | | N-terminal signal peptide from mGluR5, followed by HA, SNAP and human MC4R | | This manuscript |
| ss-HA-SNAP-mGluR2 | | Used as template for ss-HA-SNAP-MC4R | | Joshua Levitz, Weill Cornell Medicine |
| MC4R | | Used as a template for MC4R constructs | | Bryan Roth (Addgene plasmid # 66430 ; http://n2t.net/addgene:66430 ; RRID:Addgene_66430) |
| cAMP Glosensor-22F | | cAMP sensor | | Promega |
| LgBiT-IP3R2-SmBiT | | $IP_3$ biosensor | | Asuka Inoue, Tohoku University *(33)* |
| MC4R-Rluc8 | | BRET | | This manuscript |
| Venus-mGs | | BRET | | Nevin Lambert, Augusta University *(35)* |
| MRAP2-3xFLAG | | Glosensor, SIM, cell surface expression | | Julien Sebag, University of Iowa |
| MRAP2-3xFLAG-G31V | | Glosensor, IP3, SIM, cell surface expression | | Caroline Gorvin, University of Birmingham (1) |
| MRAP2-3xFLAG-P32L | | Glosensor, IP3, SIM, cell surface expression | | Caroline Gorvin, University of Birmingham (1) |
| MRAP2-3xFLAG-F62C | | Glosensor, IP3, SIM, cell surface expression | | Caroline Gorvin, University of Birmingham (1) |
| MRAP2-3xFLAG-N88Y | | Glosensor, IP3, SIM, cell surface expression | | Caroline Gorvin, University of Birmingham (1) |
| MRAP2-3xFLAG-V91A | | Glosensor, IP3, SIM, cell surface expression | | Caroline Gorvin, University of Birmingham (1) |
| MRAP2-3xFLAG-R113G | | Glosensor, IP3, SIM, cell surface expression | | Caroline Gorvin, University of Birmingham (1) |
| MRAP2-3xFLAG-S114A | | Glosensor, IP3, SIM, cell surface expression | | Caroline Gorvin, University of Birmingham (1) |
| MRAP2-3xFLAG-L115V | | Glosensor, IP3, SIM, cell surface expression | | Caroline Gorvin, University of Birmingham (1) |
| MRAP2-3xFLAG-N121S | | Glosensor, IP3, SIM, cell surface expression | | Caroline Gorvin, University of Birmingham (1) |
| MRAP2-3xFLAG-R125C | | Glosensor, IP3, SIM, cell surface expression | | Caroline Gorvin, University of Birmingham (1) |
| MRAP2-3xFLAG-H133Y | | Glosensor, IP3, SIM, cell surface expression | | Caroline Gorvin, University of Birmingham (1) |
| MRAP2-3xFLAG-T193A | | Glosensor, IP3, SIM, cell surface expression | | Caroline Gorvin, University of Birmingham (1) |
| MRAP2-3xFLAG-K42A | | Glosensor, IP3, SIM, cell surface expression | | Caroline Gorvin, University of Birmingham (1) |
| MRAP2-3xFLAG-L64A | | Glosensor, IP3, SIM, cell surface expression | | Caroline Gorvin, University of Birmingham (1) |
| MRAP2-3xFLAG-T68A | | Glosensor, IP3, SIM, cell surface expression | | Caroline Gorvin, University of Birmingham (1) |

### Table S2 Effects of MRAP2 on the predicted protein structure

| Variant | Model | Rank 1 | Rank 2 | Rank 3 | Rank 4 | Rank 5 |
|---|---|---|---|---|---|---|
| **G31V** | Monomer | E29 | None | None | E29, V33 | None |
| | Dimer | E29, V33 | Not feasible | E29, V33 | S34 | E29, V33 |
| **P32L** | Monomer | S34 | None | S34 | S34 | S34 |
| | Dimer | S34 | Not feasible | S34 | E29, V33 | S34 |
| **F62C** | Monomer | I58, L66 | I58, L66 | I58, L66 | I58, L66 | I58, L66 |
| | Dimer | I58, L66 | Not feasible | I58, L66 | I48, F59, L66. Mutant forms new contact with Ile58. | L66, N88 |
| **N88Y** | Monomer | S89 | None | R86 | M87 | None |
| | Dimer | None | Not feasible | None | None | None Mutant forms new contact with Arg86. |
| **V91A** | Monomer | None | None | None | None | None |
| | Dimer | F94 | Not feasible | S89 | D93 | D93 |
| **R113G** | Monomer | S114. Mutant loses contact. | None | None | None | None |
| | Dimer | None | Not feasible | None | E111, L115. Mutant loses Leu115 contact. | E111 |
| **S114A** | Monomer | R113. Mutant loses contact. | H117 | C118 | None | None |
| | Dimer | None | Not feasible | None | None | None |
| **L115V** | Monomer | None | None | Y119 | None | None |
| | Dimer | C118 | Not feasible | C118 | R113, C118 | H117 |
| **N121S** | Monomer | None | None | H117, R125. Mutant forms new contact with Cys118. | None | None |
| | Dimer | H117 | Not feasible | H117 Mutant forms new contact with Cys118. | C118, E124 Mutant forms new contact with Cys118. | None |
| **R125C** | Monomer | None | E122 | N121, A129 | None | None |
| | Dimer | E122 | Not feasible | None | E122, R128 | None |
| **H133Y** | Monomer | None. Mutant forms new contact with E134. | None | None | None | None |
| | Dimer | None | Not feasible | None | None. Mutant forms contact with T135. | None |
| **T193A** | Monomer | None | None | E129. Mutant loses contact. | None | None |
| | Dimer | L191 | Not feasible | None | None | None |

Table shows interactions in wild-type residues in black and the effect of the mutant residue on contacts in red. Four of the predicted MRAP2 monomers had α-helical structures in addition to the transmembrane helix. Models 2-5 had an α-helix comprising P154-M162, while models 3 and 5 had another α-helix between L115-M162 and Y119-R128, respectively. In the dimer structures, the F62 residue faces into the dimer interface in model 3 and 5, while it faces away from the dimer interface in models 1 and 4. Model 2 of the homodimer was rejected as it is not a feasible structure. Models 1, 3 and 5 have an additional α-helix between S114 and A129.

1 Jamaluddin, A., Wyatt, R.A., Lee, J., Dowsett, G., Tadross, J.A., Broichhagen, J., Yeo, G.S.H., Levitz, J. and Gorvin, C.M. (2024) The MRAP2 accessory protein directly interacts with melanocortin-3 receptor to enhance signaling. *bioRxiv*, in press., 2024.2011.2006.622243.
